# Phylogenetic position and taxonomic rearrangement of *Davidina* (Lepidoptera, Nymphalidae), an enigmatic butterfly genus new for Europe and America

**DOI:** 10.1101/2020.06.25.171256

**Authors:** Vladimir A. Lukhtanov, Vladimir V. Dubatolov

## Abstract

*Davidina*, an enigmatic butterfly genus described from China in the 19^th^ century, has been long time considered a member of the family Pieridae due to its pierid-like wing pattern. In the 20th century, it was transferred to the family Satyridae (now subfamily Satyrinae of Nymphalidae) based on analysis of genitalia structure and placed next to the species-rich genus *Oeneis* (subtribe Satyrina), being separated from the latter by supposed differences in wing venation. Here we conducted phylogenetic and taxonomic study of the subtribe Satyrina using analysis of molecular and morphological characters. We show that the genus *Oeneis* is not monophyletic, and consists of two non-sister, genetically diverged and morphologically differentiated groups (*Oeneis* s.s. and *Protoeneis*). We also demonstrate that *Davidina* is closely related to *Protoeneis*, not to *Oeneis* s.s. To avoid the discovered non-monophyly and morphological heterogeneity, several species should be extracted from *Oeneis* and transferred to the genus *Davidina*. As a consequence, we conclude that the name *Protoeneis* Gorbunov, 2001 is congeneric with *Davidina* Oberthür, 1879. We also conclude that *Davidina* is not a monotypic Chinese endemic genus as it has been previously supposed, but is composed of nine species and has a broad distribution area in the Holarctic region including Europe and America.

## INTRODUCTION

In recent years, significant progress has been made in the study of the phylogenetic relationships and taxonomic structure of butterflies (Lepidoptera, Papilionoidea) in general (Wahlberg *et al*., 2005, 2014; Espeland *et al*., 2015, 2018; Sahoo *et al*., 2016; Mitter *et al*., 2017; Seraphim *et al*., 2018; Toussaint *et al*., 2018; Zhang *et al*. 2019; Allio *et al*., 2020; Wiemers *et al*., 2020) and the species-rich family Nymphalidae in particular (Wahlberg *et al*., 2003; 2009; Leneveu *et al*., 2009; Peña *et al*., 2015; De Moya *et al.*, 2017; Kodandaramaiah *et al*., 2018; Zhou *et al*., 2020).

Within the Nymphalidae, the subtribe Satyrina has been revealed as a highly supported clade sister to the subtribe Melanargiina (Peña *et al*., 2011). According to the molecular data of Peña *et al.* (2011) and Yang & Zhang (2015), the subtribe Satyrina includes the genera *Arethusana* de Lesse, *Berberia* de Lesse, *Brintesia* Fruhstorfer, *Chazara* Moore, *Davidina* Oberthür, *Hipparchia* Fabricius, *Karanasa* Moore, *Minois* Hübner, *Neominois* Scudder, *Oeneis* Hübner, *Paroeneis* Moore, *Pseudochazara* de Lesse and *Satyrus* Latreille. This selection of genera corresponds nearly exactly to the tribe Satyrini sensu Miller, 1968 recovered on base of morphological analysis (Miller, 1968). According to Miller (1968), the following additional genera should be included in this group: *Aulocera* Butler, *Chionobas* Boisduval, *Eumenis* Hübner, *Kanetisa* Moore, *Neohipparchia* de Lesse, *Philareta* Moore and *Pseudotergumia* Agenjo.

This is a species-rich group of butterflies that includes about 200 described species (Lukhtanov, 2006). The subtribe has nearly pure Holarctic distribution, with only few species found in the Oriental region. The most speciose genera of the subtribe, such as *Chazara, Pseudochazara* and *Hipparchia* dominate in arid and steppe-like biotopes of the Palaearctic region. The species of the genus *Karanasa* inhabit alpine biotopes of the Central Asia. The genus *Oeneis* is the cold-dwelling group inhabiting taiga and tundra-like regions of the Holarctic region (Bogdanov *et al.*, 1997).

The subtribe Satyrina has been a target of several molecular phylogenetic studies (Peña *et al*., 2006, 2011, Kleckova *et al*., 2015; Yang & Zhang, 2015; Zhou *et al*., 2020). However, no one of these studies has been based on the complete sampling of the genera, and this circumstance negatively affected the reliability of the conclusions and reconstructions.

In particular, the position of *Davidina*, one of the most mysterious genera of butterflies, remains unclear. This genus is considered monotypic, being represented by a single species *Davidina armandi* Oberthür (Nakatani & Tera, 2012). It has an extremely unusual appearance for nymphalids. Its most peculiar feature is a pure white wing color in females, and whitish wing color, with more or less dark scaling in males, in combination with strongly reduced wing markings which are represented only by longitudinal dark streaks along the veins and in the intervenal spaces (Fig. 1). Similar external appearance and pigmentation are known to exist in the Palaearctic butterflies of the families Pieridae (e.g. in *Aporia* Hübner) and Papilionidae (*Parnassius subbendorfii* Ménétriès, *P. glacialis* Butler) and in the Neotropical metalmark butterfly *Styx infernalis* Staudinger (Riodinidae).

**Figure 1.**
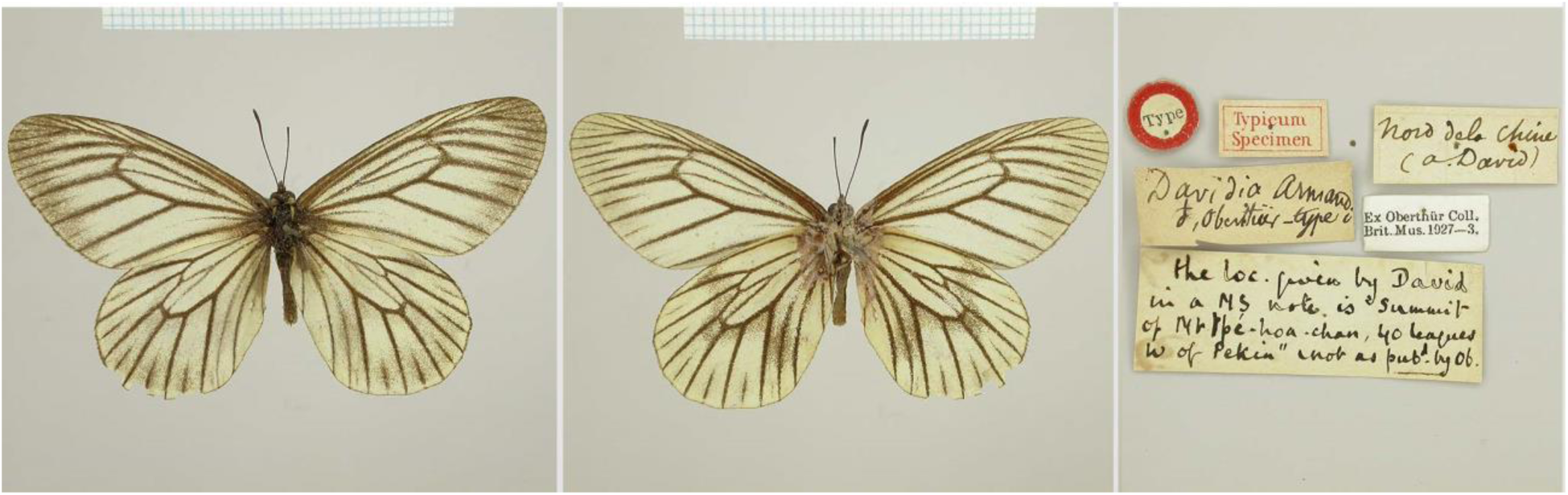
Holotype of *Davidina armandi*, upperside (left), underside (center) and labels (right). © Trustees of the Natural History Museum London, reproduced with permission. Photo: V. Lukhtanov.

There has been a much taxonomic confusion around the butterfly genus *Davidina*. The genus was described by famous French lepidopterist Charles Oberthür on base of a single male collected in China by even more famous French person Armand David (more known as abbe David). The genus was initially placed within the family Papilionidae (Oberthür, 1879). Schatz and Röber (1892) considered it as a connecting link between Papilionidae and Pieridae. Leech (1893-1894) placed *Davidina* in the family Pieridae between the genera *Aporia* and *Appias* (Leech, 1983-1984). Only in 1912 Verity (1912: page XXIV, footnote) supposed that *Davidina* is a member of the family Satyridae (now subfamily Satyrinae of the family Nymphalidae), an opinion that was confirmed by Kuznezov (1930) through thorough analysis of male and female genitalia.

After the Kuznezov’s findings, *Davidina* was accepted by entomologists as a distinct genus within the subtribe Satyrina (Miller, 1968, Lukhtanov & Eitschberger, 2000). The paper by Yang and Zhang (2015) represents the first attempt to study this genus using molecular markers; however, it did not reveal its real taxonomic position as only mitochondrial markers (and no nuclear genes) were sequenced for *Davidina*, and closely related genera *Oeneis, Karanasa* and *Neominois* were not included in the analysis.

Here, in order to reveal the phylogenetic position and taxonomic composition of this genus, we analyzed the wing venation in *Davidina* and *Oeneis*, studied the male genitalia in *Davidina* and the closely related genera *Oeneis, Neominois, Karanasa* and *Paroeneis* and conducted molecular phylogenetic analyses of the subtribe Satyrina using three data sets: (1) mitochondrial gene *COI* barcodes, (2) concatenation of nuclear genes *wingless*+*RpS5*+*GAPDH* and mitochondrial gene *COI*, and (3) nuclear gene *EF1-a*.

## MATERIAL AND METHODS

### Sampling

We studied the following specimens of *D. armandi*: one female from the Siberian Zoological Museum, Novosibirsk (used for molecular studies and analysis of wing venation), four historical samples (three males and one female) from Zoological Institute of the Russian Academy of Sciences, St. Petersburg (for analysis of male genitalia and wing venation); three historical samples (including holotype of *Davidina armandi*, Fig. 1) from Natural History Museum London (for analysis of wing venation); twelve samples from the McGuire Center for Lepidoptera and Biodiversity, University of Florida (for molecular studies and analysis of wing venation).

For molecular and morphological analyses, we also used the samples of the genera *Arethusana, Aulocera, Brintesia, Chazara, Hipparchia, Karanasa, Melanargia, Minois, Neominois, Oeneis, Paroeneis, Pseudochazara* and *Satyrus* preserved in Zoological Institute of the Russian Academy of Sciences. The accession numbers of these samples are presented in Figures 2-4.

**Figure 2.**
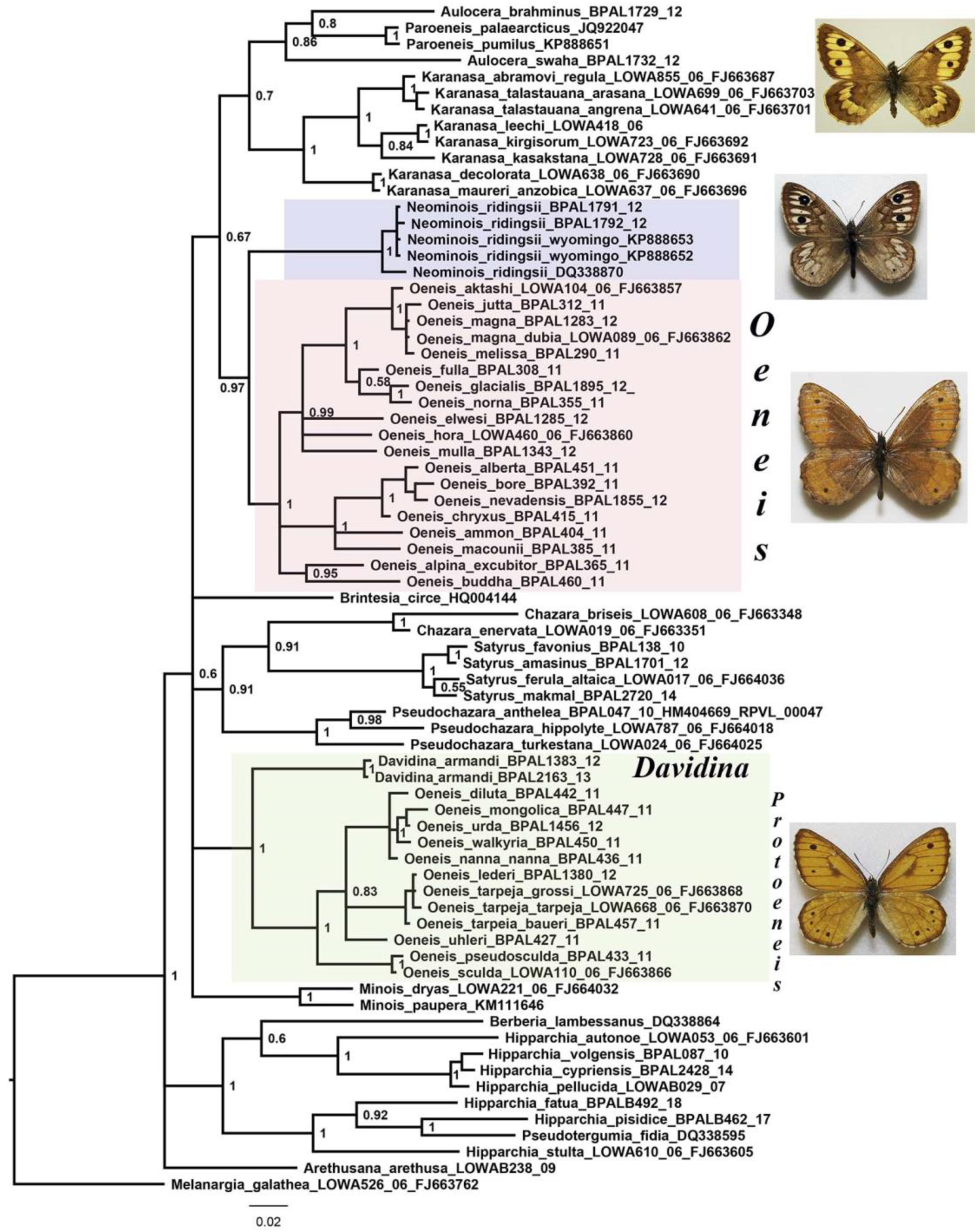
Phylogenetic tree from BI analysis of 72 taxa of the subtribe Satyrina based on mitochondrial *COI* barcodes. Posterior probabilities are indicated at the nodes.

**Figure 3.**
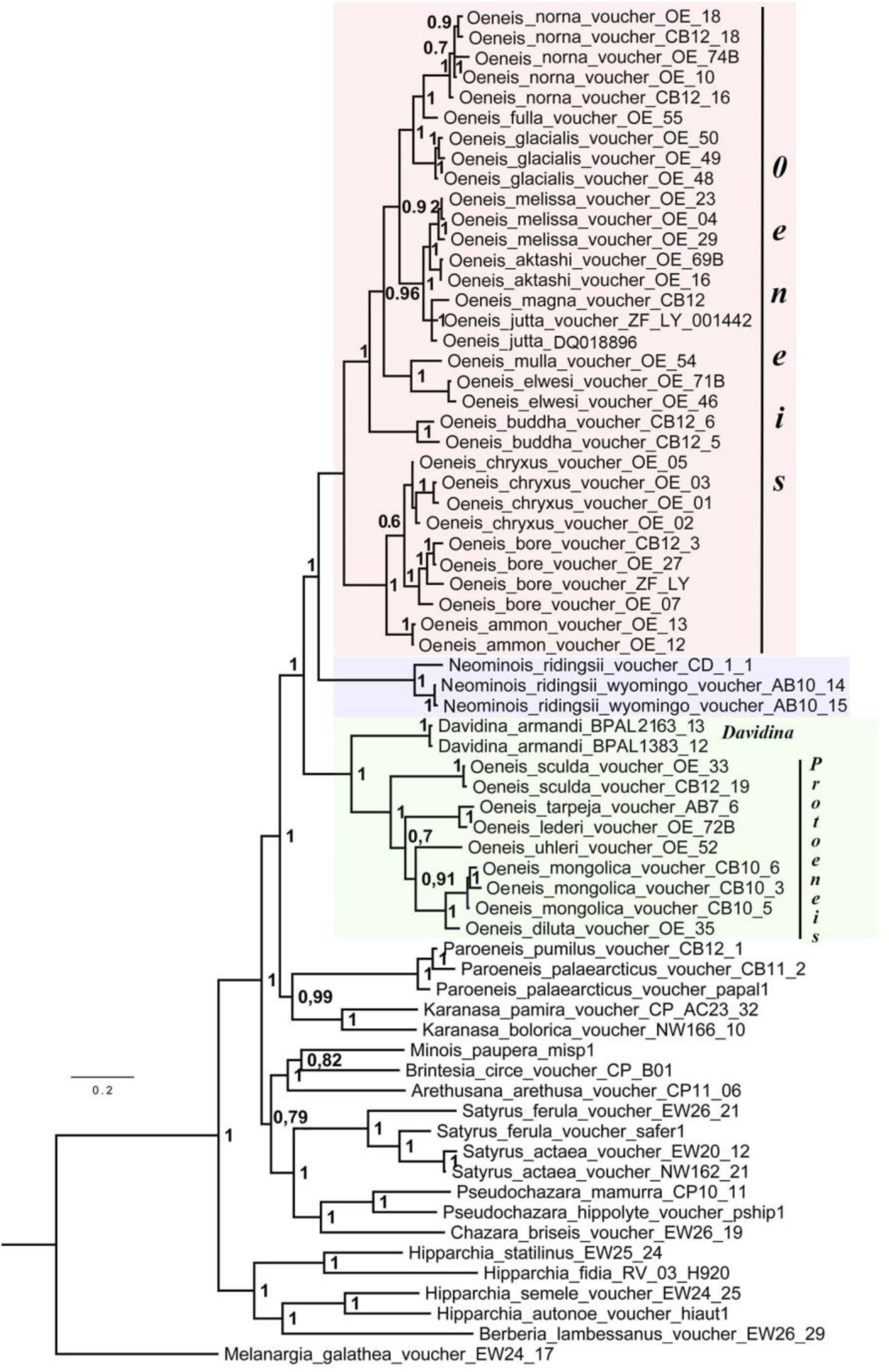
Phylogenetic tree from BI analysis of 66 samples of the subtribe Satyrina based on *COI+wingless*+*RpS5*+*GAPDH* data set. Posterior probabilities are indicated at the nodes.

**Figure 4.**
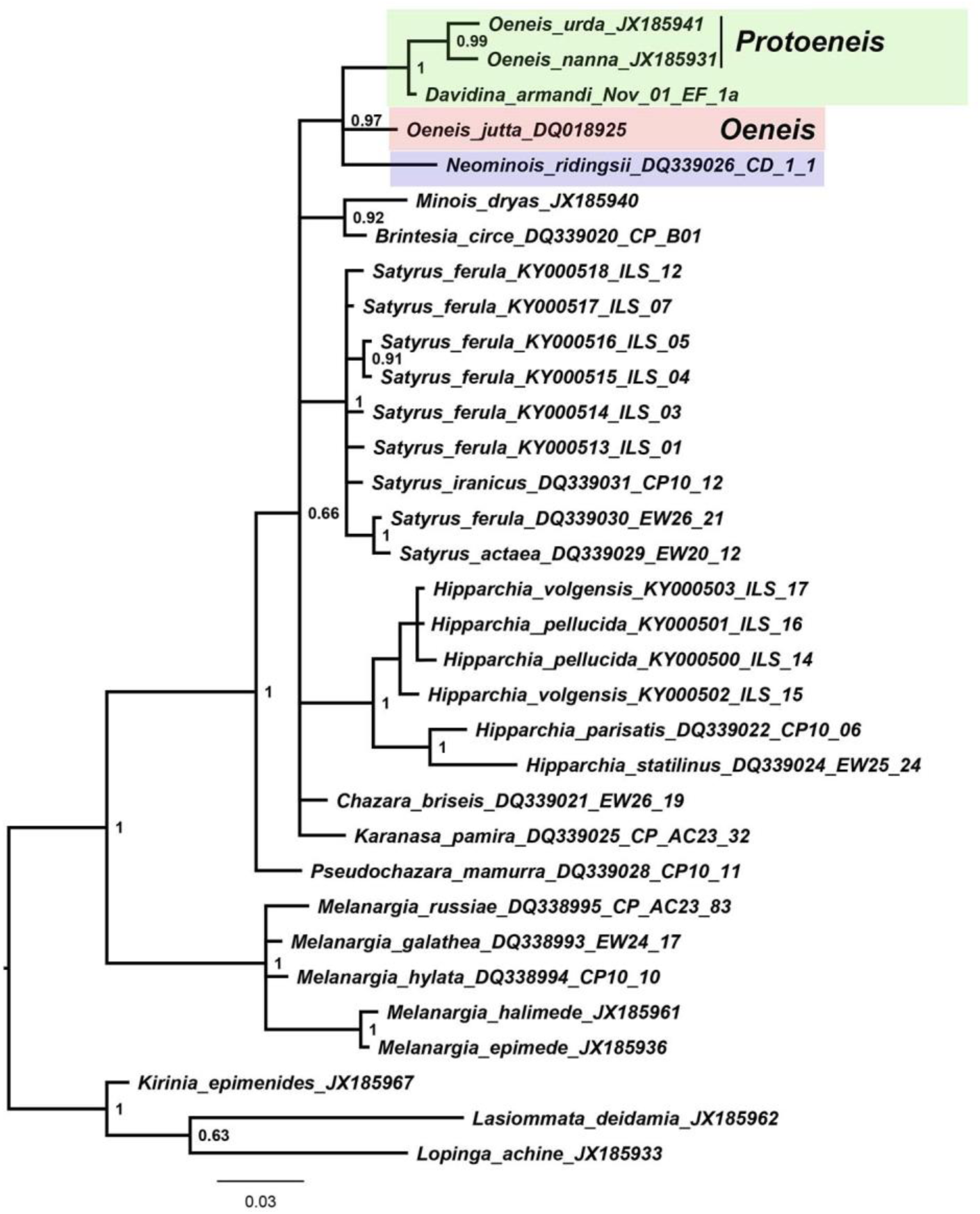
Phylogenetic tree from BI analysis of 30 samples of the subtribe Satyrina and 3 outgroup samples based on the nuclear gene *EF1a*. Posterior probabilities are indicated at the nodes.

### DNA studies

Standard *COI* barcodes (partial sequences of the *cytochrome c oxidase subunit I* gene) were obtained from a single leg of two *D. armandi* and 37 samples of other representatives of the subtribe Satyrina at the Canadian Centre for DNA Barcoding (CCDB, Biodiversity Institute of Ontario, University of Guelph) using their standard high-throughput protocol described by deWaard *et al.* (2008). The pictures, and collection data of these specimens are deposited and can be freely downloaded at the BOLD Public Data Portal (http://www.boldsystems.org/index.php/databases). The sequence of the nuclear gene *EF1-a* was obtained using primers and protocols described in Stradomsky *et al*. (2016). The pictures and collection data of this specimen are deposited and can be freely downloaded at Siberian Zoological Museum, Novosibirsk (http://szmn.eco.nsc.ru/picts/butterfly/Satyridae/Davidina_armandi.htm).

We constructed phylogramms using three data sets: (1) *COI* barcodes, (2) concatenation of nuclear genes *wingless*+*RpS5*+*GAPDH* and mitochondrial *COI* barcode gene, and (3) nuclear gene *EF1-a*. For these analyses we used our own sequences as well as published sequences extracted from GenBank (Peña *et al*., 2006, 2011, Lukhtanov *et al*., 2009; Dincă *et al*., 2011; Wan *et al*., 2013; Kleckova *et al*., 2015, Yang & Zhang, 2015; Stradomsky *et al*., 2016). The GenBank/BOLD/museum accession numbers of the analyzed sequences are presented in Figs 2-4. *Melanargia galathea* was selected as outgroup in accordance with the published data (Peña *et al*., 2011).

Sequences were aligned using BioEdit software (Hall, 1999) and edited manually. The *COI* alignment length was 655 bp (236 variable and 197 parsimony informative sites), the *wingless*+*RpS5*+*GAPDH*+ *COI* alignment length was 3195 bp (898 variable and 664 parsimony informative sites) The *EF1-a* alignment length was 493 bp (122 variable and 101 parsimony informative sites).

Nucleotide substitution models for each dataset was estimated based on the Bayesian Information Criterion using jModeltest, version 2 (Darriba *et al*., 2012, Guindon & Gascuel, 2003). The best fitting models were as follows: GTR + G + I for *COI* and for *wingless*+*RpS5*+*GAPDH*+ *COI;* and K2 + G for *EF1-a*.

The Bayesian analyses (Bayes Inference, BI) were performed using the program MrBayes 3.2 (Ronquist *et al*., 2012): with default settings as suggested by Mesquite (Maddison & Maddison, 2015): (lset nst=6 rates=invgamma; burnin=0.25) for the *COI* and for *wingless*+*RpS5*+*GAPDH*+ *COI* data sets and with the settings (lset nst= rates=gamma; burnin=0.25) for the *EF1-a* alignment. Two runs of 10,000,000 generations with four chains (one cold and three heated) were performed. The consensus of the obtained trees was visualized using FigTree 1.3.1 (http://tree.bio.ed.ac.uk/software/figtree/).

### Morphological analysis

For genitalia preparation adult abdomens were soaked in hot (90°C) 10% KOH for 3-10 min. Then they were transferred to water, the genitalia were carefully extracted and macerated under a stereo-microscope with the help of a pair of preparation needles or with help of a needle and a watchmaker’s tweezer. Once cleansed of all unwanted elements they were transferred and stored in glycerine. Cleansed genitalia armatures were handled, studied and photographed while immersed in glycerine, free from pressure due to mounting, and therefore free from the ensuing distortion. Genitalia photographs were taken with Leica M205C binocular microscope equipped with Leica DFC495 digital camera, and processed using the Leica Application Suite, version 4.5.0 software. In order to evaluate intraspecific variation in genitalia structure, three males of *Davidina armandi*, five males of *Oeneis* (*Protoeneis*) *nanna*, five males of *Oeneis* (*Oeneis*) *norna*, four males of *Neominois ridingsii*, three males of *Paroeneis palearcticus* and five males of *Karanasa kirgisorum* were studied.

Butterfly photographs were taken with Nikon D810 digital camera equipped with Nikon AF-S Micro Nikkor 105 mm lens.

### Phylogenetic analysis of the morphological characters

Twelve characters of the male genitalia were selected for the analysis (Table 1). This selection was based on comparative analysis of male genitalia in the genera and subgenera of the subtribe Satyrina. For the genera and subgenera *Davidina, Oeneis* (*Oeneis*), *Oeneis* (*Protoeneis*), *Neominois, Paroeneis* and *Karanasa*, we used our original data. For the genera *Arethusana, Aulocera, Brintesia, Chazara, Hipparchia, Minois, Pseudochazara* and *Satyrus* we used published data (Nekrutenko, 1985; Chou, 1998; Jakšić, 1998; Sharma & Rose, 2014).

**Table 1.**
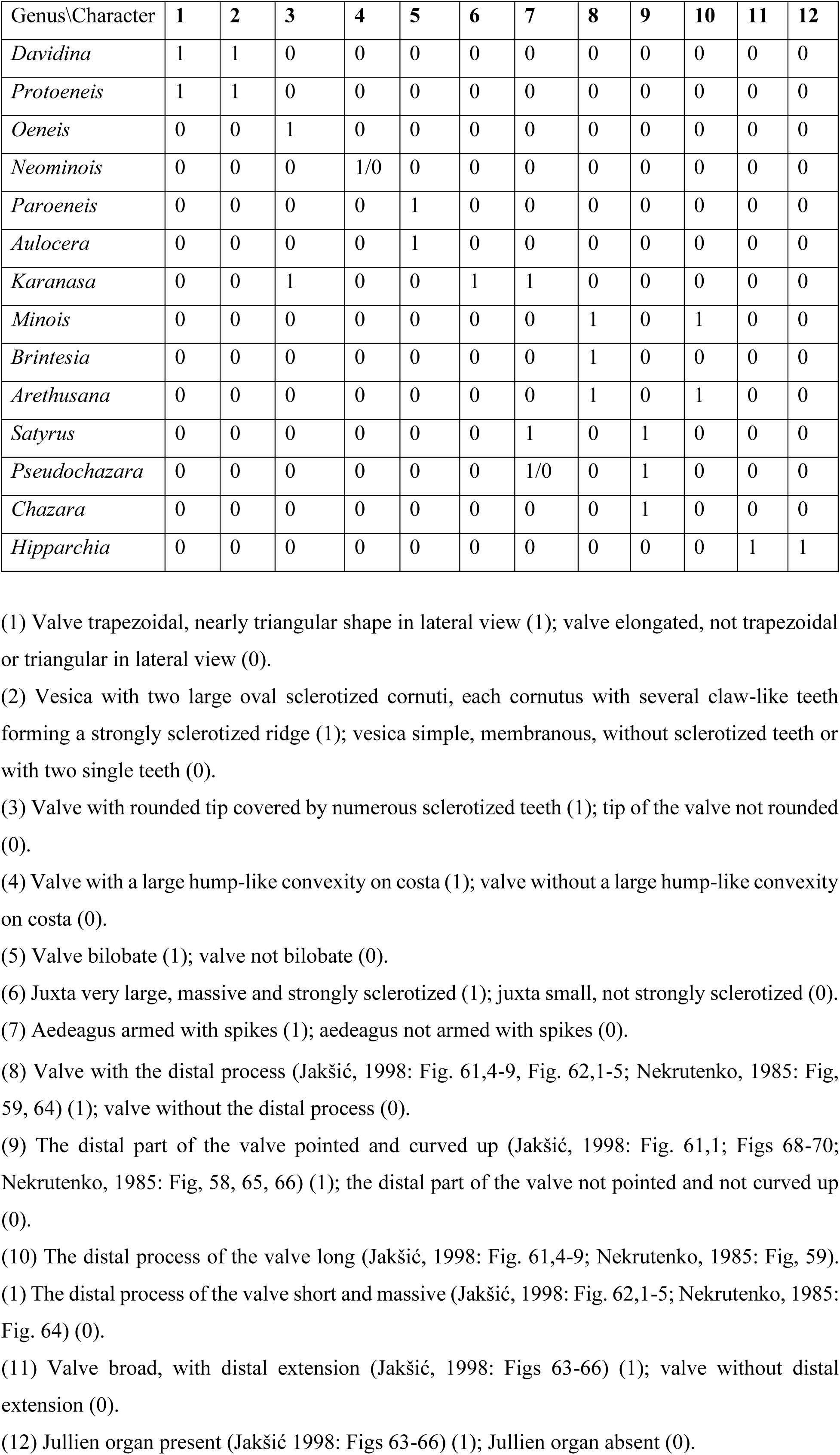
Matrix of morphological characters

All the characters were binary and equally weighted. The variable characters 4 in *Neominois* and 7 in *Pseudochazara* (Table 1) were coded as “?”. Maximum parsimony phylogenetic analyses (MP) were carried out in PAUP* (Swofford, 2002). Analyses were conducted holding up to 10000 trees and searching for 100 replicates under tree-bisection-and-reconnection (TBR) branch swapping. We also tested for nodal support using the Bootstrap method with 1000 replicates. Bayesian analyses (BI) of the binary characters we performed using the program MrBayes 3.2 (Ronquist *et al*., 2012) and the following command block:

~~~
begin mrbayes;
    set autoclose=yes nowarn=yes;
    unlink shape=(all) pinvar=(all) statefreq=(all) revmat=(all);
    prset applyto=(all) ratepr=variable;
    mcmcp ngen= 10000000 relburnin=yes burninfrac=0.25 printfreq=10000
samplefreq=10000 nchains=4 savebrlens=yes;
    mcmc;
    sumt;
end;
~~~

Two runs of 10 000 000 generations with four chains (one cold and three heated) were performed. Chains were sampled every 10000 generations.

The genus *Hipparchia* is known to have a well supported basal position within the subtribe Satyrina (Peña *et al*., 2011) and was used to root the rest of the Satyrina.

## RESULTS

### Phylogenetic analyses

#### Phylogenetic relationships based on mitochondrial gene COI

Analysis based on mitochondrial gene *COI* (Fig. 2) revealed several highly supported monophyletic lineages within the subtribe Satyrina. The majority of these lineages correspond to the traditional genera *Paroeneis, Karanasa, Neominois, Chazara, Satyrus, Pseudochazara* and *Minois*. Interestingly, *Oeneis* was revealed as a clearly non-monophyletic assemblage consisting of two not closely related clusters. One cluster was presented by the species of the subgenus *Oeneis* (*Oeneis*), the other clusters was represented by species of the subgenus *Oeneis* (*Protoeneis*).

The genus *Neominois* was found as a sister to *Oeneis* (*Oeneis*), and the genus *Davidina* was found as a sister to *Oeneis* (*Protoeneis*). The monophyly of the clades [*Oeneis* (*Oeneis*)+*Neominois*] and [*Oeneis* (*Protoeneis*)+*Davidina*] was highly supported. The phylogenetic relationships between other genera are unresolved or have low support on the *COI* tree, most likely because of insufficient phylogenetic power of this marker.

#### Phylogenetic relationships based on concatenation of mitochondrial gene COI and nuclear genes wingless+RpS5+GAPDH

Concatenation of the nuclear genes *wingless+RpS5+GAPDH* with mitochondrial *COI* barcodes (for *Davidina* only mitochondrial *COI* barcodes were available) resulted in the phylogenetic reconstruction in which the majority of the main nodes were highly supported (Fig. 3).

The target genera of our research, *Oeneis, Neominois, Davidina, Karanasa* and *Paroeneis*, were found to constitute a monophyletic group. Within this group, the genus *Oeneis* was found as a non-monopyletic assemblage. The subgenus *Oeneis* (*Oeneis*) was found as a sister to the genus *Neominois*, and the subgenus *Oeneis* (*Protoeneis*) was found as a sister to the genus *Davidina*.

#### Phylogenetic relationships based on nuclear gene EF1a

Although this analysis was based on a less number of taxa, it was completely consistent with the conclusions based on *COI* and *COI+wingless*+*RpS5*+*CAPDH* data sets. The representatives of the thee target genera of our research, *Oeneis, Neominois* and *Davidina* were found to constitute a highly supported monophyletic group (Fig. 4).

Within this group, the genus *Oeneis* was found as a non-monopyletic assemblage. *Davidina armandi* was found as a sister to a group *O. nanna* + *O. urda* (subgenus *Protoeneis*) supporting the close relatedness of *Davidina* and *Protoeneis*.

### Male genitalia

#### *Davidina armandi* (type species of *Davidina*) (Figs 5a, 6a)

Valve angulate in lateral view, trapezoidal; the apical part of the valve acutely pointed; there are no sclerotized teeth on the valve. Tegumen massive. Uncus very massive and broad. Subuncii thin and pointed, they have approximately the same thickness in entire length. Juxta small, not strongly sclerotized. Aedeagus long. Vesica with two large oval cornuti, each cornutus with several (7-9) claw-like teeth forming a strongly sclerotized ridge.

**Figure 5.**
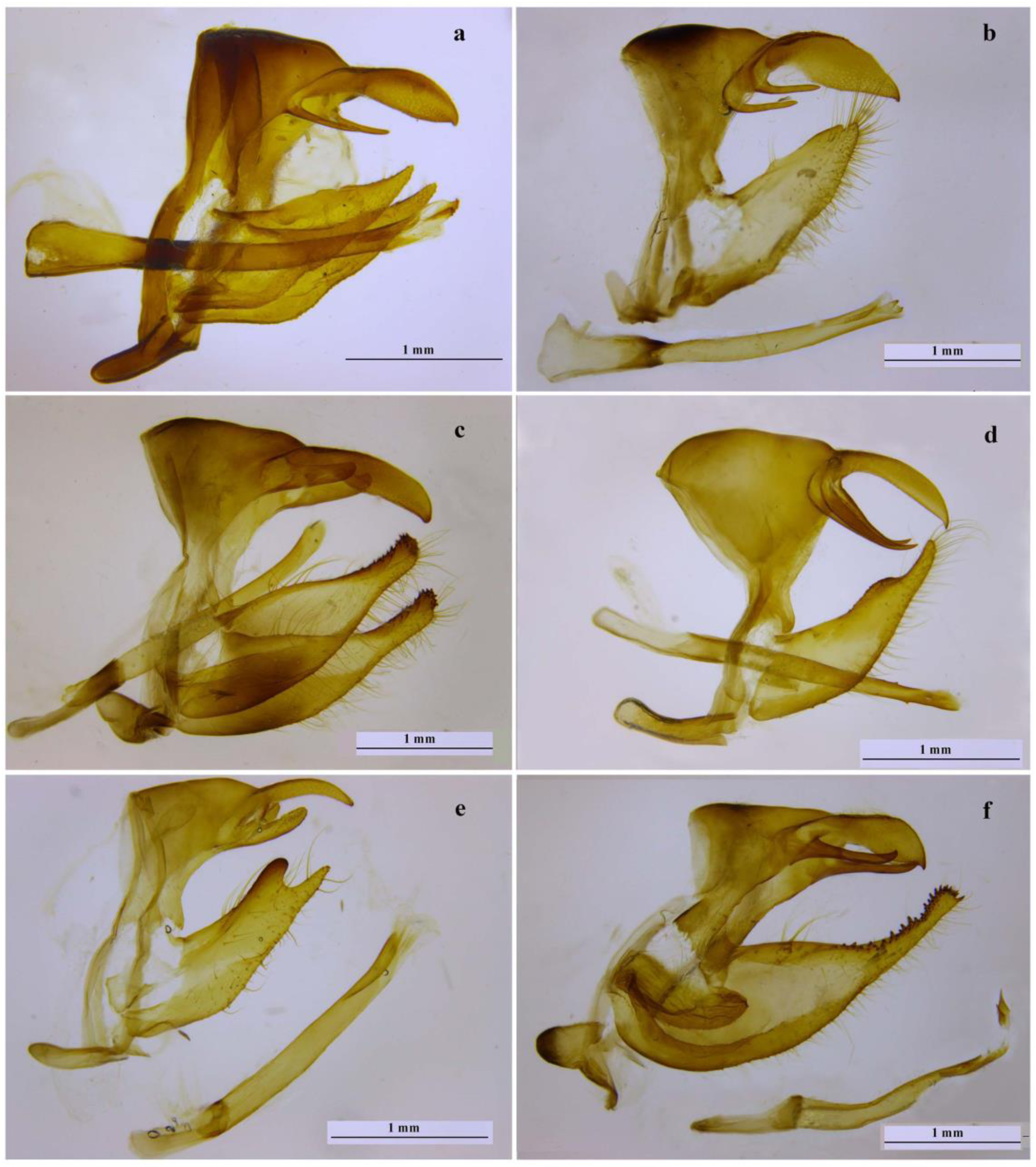
Male genitalia, lateral view. (a) *Davidina armandi*, China, Nei Mongol, Inn-Shan Mts; (b) *Oeneis* (*Protoeneis*) *nanna* Ménétries; Russia, Buryatia, Sosnovka; right valve is not shown, aedeagus is separated; (c) *Oeneis* (*Oeneis*) *norna* Thunberg, Russia, Altai; (d); *Neominois ridingsii* Edwards, USA, Colorado; left valve is not shown; (e) *Paroeneis palaearcticus* Staudinger, China, Xinjian/Qinghai, Altyn-Tag, right valve is not shown, aedeagus is separated; (f) *Karanasa kirgisorum* Avinov et Sweadner, Kazakhstan, Kirgyzsky Mts, left valve is not shown, aedeagus is separated.

**Figure 6.**
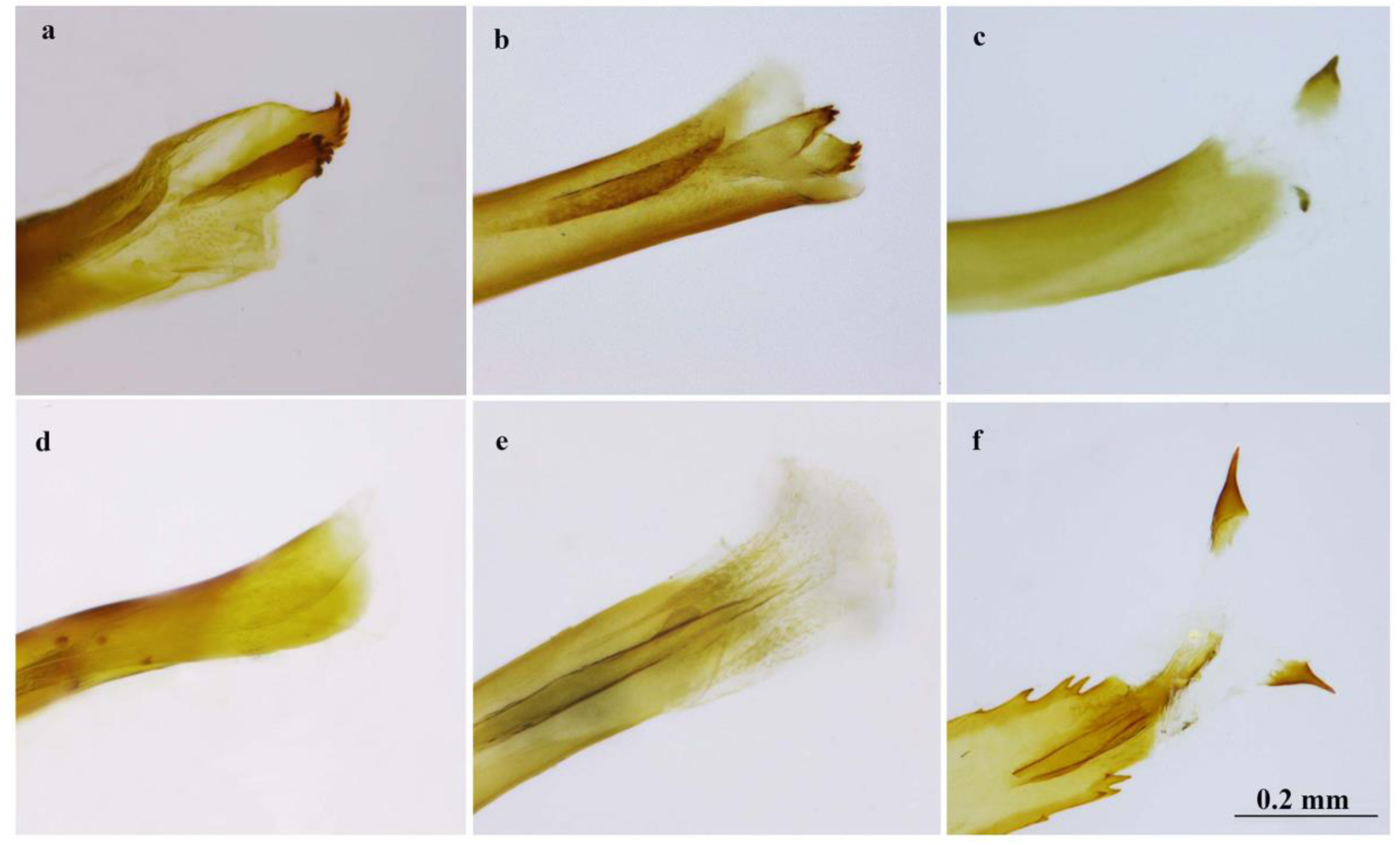
Apical part of aedeagus (a) *Davidina armandi*, China, Nei Mongol,Inn-Shan Mts; (b) *Oeneis* (*Protoeneis*) *nanna*, Russia, Buryatia, Sosnovka; (c) *Oeneis* (*Oeneis*) *norna*, Russia, Altai; (d) *Neominois ridingsii*, USA, Colorado; (e) *Paroeneis palaearcticus*, China, Xinjian/Qinghai Altyn-Tag, (f) *Karanasa kirgisorum*, Kazakhstan, Kirgyzsky Mts.

#### *Oeneis nanna* Ménétries (type species of *Protoeneis*) (Fig. 5b, 6b)

Valve angulate in lateral view, trapezoidal; the apical part of the valve not pointed; there are no sclerotized teeth on the valve. Tegumen massive. Uncus very massive and broad. Subuncii thin and pointed, they have approximately the same thickness in entire length. Juxta small, not strongly sclerotized. Aedeagus long. Vesica with two large oval sclerotized cornuti, each cornutus with several (4-6) claw-like teeth forming a strongly sclerotized ridge.

#### *Oeneis norna* Thunberg (type species of *Oeneis* s.s) (Fig. 5c, 6c)

Valve elongated in lateral view, not trapezoidal, with rounded tip; there are sclerotized teeth on the dorsal part of valve near the tip. Tegumen massive. Uncus not massive. Subuncii broad and not pointed, they have approximately the same thickness in entire length. Juxta small, not strongly sclerotized. Aedeagus long. Vesica entirely different from *Davidina* and *Oeneis* (*Protoeneis*) with two tooth-like, sharply terminated cornuti.

#### *Neominois_ridingsii* Edwards (type species of *Neominois*) (Fig. 5d, 6d)

Valve with a specific hump-like convexity on costa; without sclerotized teeth. Tegumen massive. Uncus not massive. Subuncii thin and pointed, they gradually thin to the top. Aedeagus long. Vesica simple, membranous, without sclerotized teeth.

#### *Paroeneis palaearcticus* Staudinger (Figs 5e, 6e)

Valve bilobate, with entirely different shape as compared with other genera. There are no sclerotized teeth on the valve. Tegumen massive. Uncus not massive. Subuncii broad and not pointed, they have approximately the same thickness in entire length. Juxta small, not strongly sclerotized. Aedeagus long. Vesica simple, membranous, without sclerotized teeth

#### *Karanasa kirgisorum* Avinov et Sweadner (Figs 5f, 6f)

Valve elongated in lateral view, not trapezoidal, with rounded tip; there are sclerotized teeth on the dorsal part of valve near the tip. Tegumen massive. Subuncii have approximately the same thickness in entire length. Juxta massive and strongly sclerotized. Aedeagus relatively short, curved and armed with spikes. Vesica with two cornuti, each possessing a single sclerotized tooth. Generally, the male genitalia in *Karanasa* are similar to those in *Oeneis* (*Oeneis*), except for the massive and strongly sclerotized juxta and the aedeagus, which is curved and armed with spikes.

Thus, the following characters of the male genitalia are unique for one or two of the studied genera and subgenera and represent potential (syn)apomorphies:

-trapezoidal, nearly triangular shape of valve in lateral view (character 1) and vesica with two large oval sclerotized cornuti, each cornutus with several claw-like teeth forming a strongly sclerotized ridge (character 2) (unique characters for *Davidina* and *Oeneis* (*Protoeneis*);
-valve distinctly narrowed in distal portion, not trapezoidal, with rounded tip (character 3) (unique character for *Oeneis* (*Oeneis*) and *Karanasa*);
-specific shape of valve with a hump-like convexity on costa (character 4). This character is found in *Neominois ridingsii* (de Lesse, 1951; Warren et al., 2008; this study) and seems to be unique for *Neominois*. However, it is not genus-specific since it is not found in the second species of *Neominois, N. carmen* Warren, Austin, Llorente, Luis & Vargas, 2008 (Warren et al., 2008);
-bilobate valve (character 5), found by us for *Paroeneis* and previously reported for *Aulocera* (de Lesse, 1951; Chou, 1998; Sharma & Rose, 2014);
-massive and strongly sclerotized juxta (character 6) and aedeagus armed with spikes (character 7) (unique character for *Karanasa*).

### Phylogenetic analysis of the morphological characters

Based on the original data on characters 1-7 and published data on characters 4, 8-12 (de Lesse, 1951; Nekrutenko, 1985, Jakšić, 1998; Warren et al., 2008), we created the following matrix of the binary characters (Table 1):

MP analysis of the morphological matrix revealed five clades that appeared in all 9 discovered MP trees. The 100% consensus of the 9 MP trees is shown in Fig. 7.

**Figure 7.**
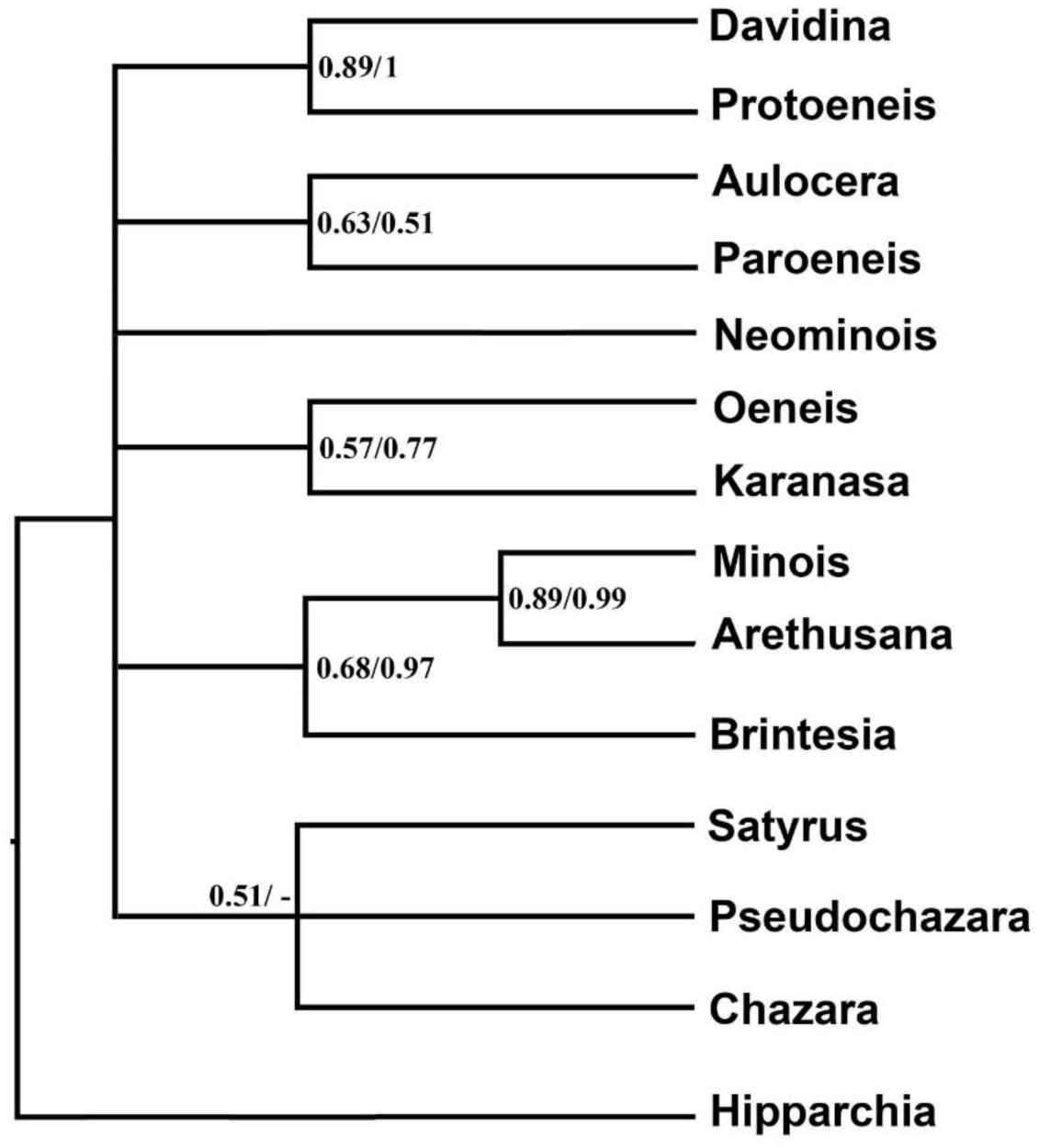
100% consensus of the 9 MP trees based on the matrix of 12 morphological characters (Table 1). BI of the morphological matrix revealed the same topology. Bootstrap support for MP/posterior probabilities for BI values are indicated at the nodes. The sign “-” indicates that the value is lower than 0.5. The genus *Hipparchia* is known to have a basal position within the subtribe Satyrina (Peña et al. 2011) and was used as outgroup for the rest of the Satyrina.

The same topology was revealed in by BI analysis of the morphological matrix revealed. Three clades, (*Davidina*+*Protoeneis*), (*Minois*+*Arethusana*) and (*Brintesia* + (*Minois*+*Arethusana*)) were highly supported in BI analysis and had a medium support in MP analysis. Three clades, (*Aulocera*+*Paroeneis*), (*Oeneis*+*Karanasa*) and (*Satyrus*+*Pseudochazara*+*Chazara*) had low support.

### Wing venation

According to Kuznetsov (1930) there is a clear distinction between *Davidina* (based on *D. armandi*, the type species of the genus) and *Oeneis* (based on *O. norna*, the type species of the genus) in the fore wing venation. First, in *Davidina* all radial branches, except R1, form a common and rather long stalk R2+3+4+5; in *Oeneis* r2 is free, i,e, R2 and R3+4+5 start independently from the same point of the discal cell. Second, in *Davidina* M1 rises at a distance from this stalk, the vein r-m1 being, thus, well developed; in *Oeneis* M1 rises from one point with R3+4+5, and r-m1 is, this, absent.

However, Kuznetsov’s observation was based on two specimens only. We analyzed 18 specimens of *D. armandi* and discovered that four types of venation are present in *Davidina*:

1. all radial branches, except R1, form a common and rather long stalk R2+3+4+5; M1 rises at a distance from this stalk, the vein r-m1 being, thus, well developed (*Davidina* type according to Kuznetsov, 1930, found in 11 samples, Fig. 8a);
2. veins R2, R3+4+5 and M1 rise from the same point on the discal cell; the vein r-m1 is absent (*Oeneis* type according to Kuznetsov, 1930, found in four samples, Fig. 8b);
3. veins R2 and R3+4+5 rise from the same point on discal cell; M1 rises at a distance from this point, the vein r-m1 being, thus, well developed (found in two samples, Fig. 8c);
4. all radial branches, except R1, form a common and rather long stalk R2+3+4+5; this stalk and M1 rise from the same point on the discal cell; the vein r-m1 is absent (found in a single sample, not shown).

**Fig. 8.**
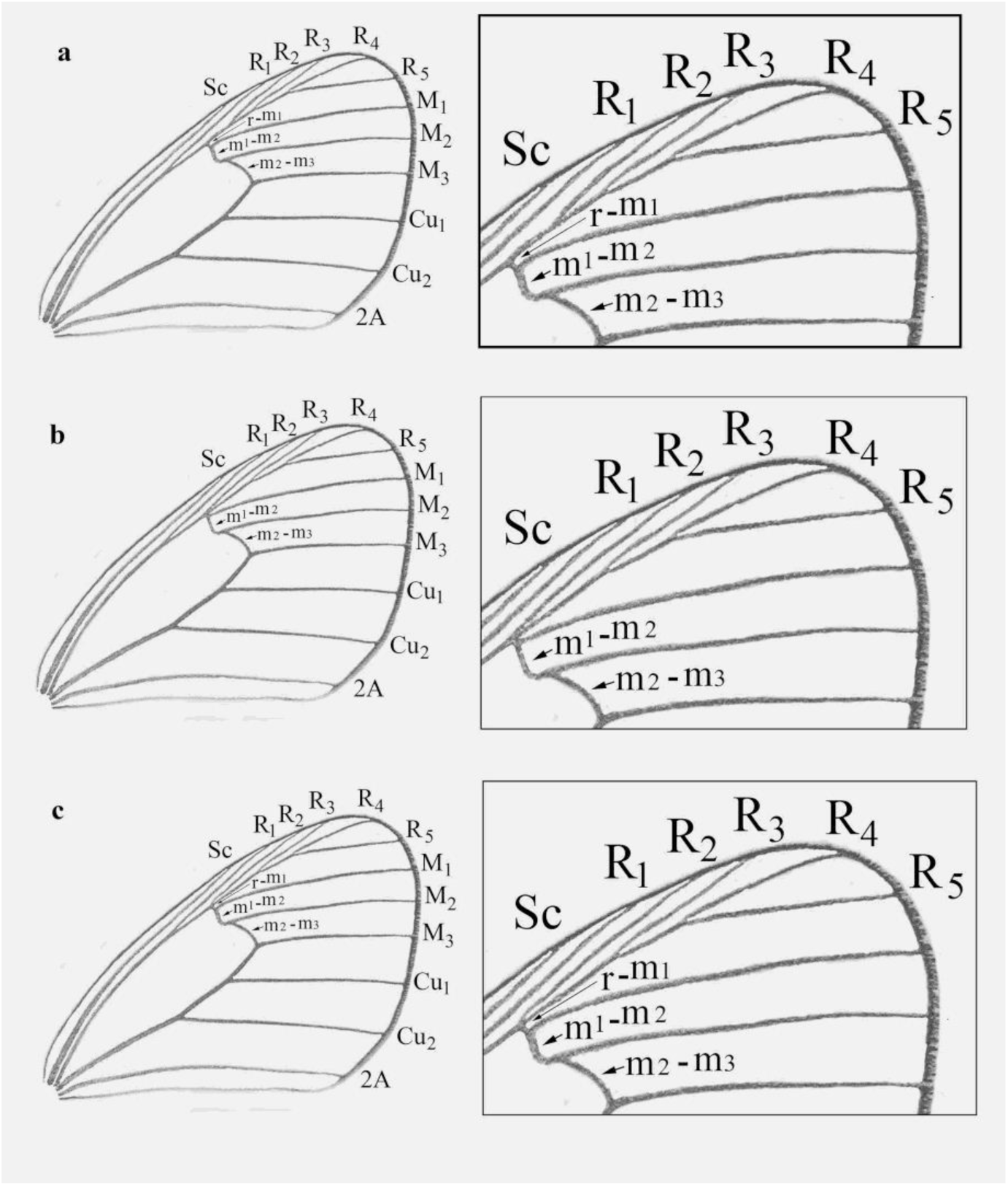
Different types of the fore wing venation in *Davidina armandi*. a) all radial branches, except R1, form a common and rather long stalk R2+3+4+5; M1 rises at a distance from this stalk; b) veins R2, R3+4+5 and M1 rise from the same point on the discal cell; the vein r-m1 is absent; c) veins R2 and R3+4+5 rise from the same point on discal cell; M1 rises at a distance from this point.

Thus, the wing venation is not stable in *D. armandi*. It is not genus-specific, but variable even on population level. The supposed differences between *Oeneis* and *Davidina* in wing venation (Kuznezov, 1930) is not confirmed.

## DISCUSSION

### The phylogeny of the subtribe Satyrina

All three molecular analyses demonstrate that (1) the genus *Oeneis* in its traditional composition (Gross, 1970; Lukhtanov 1985; Lukhtanov & Eitschberger, 2000, 2001) is not monophyletic. In fact, this is an assemblage consisting of two non-sister lineages, *Oeneis* (*Oeneis*) and *Protoeneis.* Additionally, all these analyses show that *Davidina* is a sister to the subgenus *Oeneis* (*Protoeneis*).

In the analyses based on *COI* and *COI+wingless*+*RpS5*+*GAPDH*, the genus *Neominois* is found in the position between *Oeneis* and [*Davidina* + *Protoeneis*], being a sister to *Oeneis* s.s. The sister relationships between *Oeneis* s.s and *Neominois* have been demonstrated in the previous molecular studies (Peña et al., 2011; Kleckova et al. 2015). Analysis based on *EF1a* did not resolve the phylogenetic position of *Neominois*, but confirmed the grouping *Oeneis* (*Oeneis*), *Oeneis* (*Protoeneis*), *Neominois* and *Davidina* in a single highly supported clade.

The butterflies of the genus *Paroeneis* have a wing pattern similar to that in *Oeneis* (*Oeneis*) and *Oeneis* (*Protoeneis*), and the wing pattern in *Karanasa* is similar to that in *Neominois*. Despite this, the genera of the pairs *Paroeneis* - *Oeneis* and *Karanasa* – *Neominois* have not been revealed as sisters. Instead, the genera *Karanasa* and *Paroeneis* are found to constitute a clade in the analysis based on *COI+wingless*+*RpS5*+*GAPDH*. In turn, the clade [*Karanasa*+*Paroeneis*] is found as a sister to [(*Oeneis*+*Neominois*)+(*Davidina*+ *Protoeneis*)].

To check the correctness of the revealed phylogenetic position of *Davidina* as a sister to *Oeneis* (*Protoeneis*), we conducted a comparative analysis of male genitalia in the group [*Karanasa*+*Paroeneis*]+[(*Oeneis*+*Neominois*)+(*Davidina*+ *Protoeneis*)]. Morphological analysis revealed two characters in the structure of genitalia, which can be interpreted as synapomorphies of *Davidina* and *Protoeneis*. These are the shape of the valve and the structure of the vesica. The latter is transformed in *Davidina* and *Protoeneis* in a complex organ with a specific system of teeth (Fig. 6a, b). The independent and parallel origin of this organ in *Davidina* and *Protoeneis* seems unlikely. The MP and BI phylogenetic analyses of the morphological characters also supported the clade (*Davidina* + *Protoeneis*).

The analyzes based on molecular data and morphology led to congruent results, although the resolution of the morphological matrix was lower. Within the subtribe Satyrina, molecular analysis revealed the following four groups of genera:

1. group of genera and subgenera close to *Oeneis* (*Oeneis, Protoeneis, Davidina, Neominois, Paroeneis, Karanasa*);
2. group of genera close to *Brintesia* (*Brintesia, Arethusana, Minois*);
3. group of genera close to *Satyrus* (*Satyrus, Chazara, Pseudochazara*);
4. *Hipparchia* (including *Berberia*).

Of these groups, the second, third and fourth groups were also identified in morphological analysis.

### Taxonomic interpretation of the revealed phylogenetic pattern

The data obtained raise the question about how the discovered monophyletic groups close to *Oeneis* (*Oeneis* sensu stricto, *Protoeneis, Neominois*, and *Davidina*) should be interpreted from the viewpoint of taxonomy. There are several ways to interpret the discovered topology in purpose of taxonomy:: (1) all these lineages represent a single genus with four subgenera, (2) there are two genera in the group, (*Davidina*+*Protoeneis*) and (*Oeneis*+*Neominois*); (3) each lineage represents a genus, so there are four genera in the group; (4) there are three genera, (*Davidina*+*Protoeneis*), *Oeneis* s.s/ and *Neominois*.

In our opinion, the merging these lineages in two, and even more so, in a single genus, seems unacceptable for the following reasons. (1) The divergence time analysis indicated that the split between the lineage leading to *Protoeneis* and the lineage (*Oeneis* + *Neominois*) occurred around 12 MYA, and the split between *Neominois* and *Oeneis* occurred around 10 MYA (Kleckova *et al*., 2015). This time far exceeds the interval (5 million), which has been proposed as one of the criteria for the separation on the genus level (Talavera *et al*., 2013). Although this interval has been proposed for representatives of another family, Lycaenidae, it was de facto recently used to delimit genera of nymphalids: and a similar age (7.5-9.1 MYA) was used to support the genus status of *Argynnis* Fabricius, *Fabriciana* Reuss and *Speyeria* Scudder (De Moya *et al*., 2017). (2) This merging will result in a highly heterogeneous group(s) from a viewpoint of morphology. No morphological synapomorphies are known for the grouping *Oeneis* s.s +*Protoeneis*+*Neominois*+*Davidina*, as well as for *Neominois*+*Davidina.* (3) Their merging will affect the issue of stability of the nomenclature. *Neominois* and *Davidina* have never been considered as members of the genus *Oeneis* (the only exception is the recent work by Klekova *et al*. (2015) where *Neominois* was included in *Oeneis*).

The four-genus system is also inappropriate because there are no reasons to differentiate *Davidina* and *Protoeneis* as distinct genera. (1) The level of genetic divergence between them is small (3.19% of the fixed differences in the *COI* barcode region), and applying the “standard” mitochondrial substitution rate 1.5-2.3 % uncorrected pairwise distance per million years (Brower, 1994; Quek *et al*., 2004) results in the divergence time less than 5 MY. (2) There is no significant difference between *Davidina* and *Protoeneis* in male genitalia. (3) We did not confirm the supposed (Kuznezov, 1930) difference between *Oeneis* and *Davidina* in wing venation. 4) At first glance, there is a significant difference between *D. armandi* and the species of the subgenus *Protoeneis* in the wing color and pattern. *Davidina armandi* looks unusual because of extreme reduction of all elements of the wing pattern, except longitudinal dark streaks along the veins and in the intervenal spaces. However, a tendency to such a reduction is observed in other species of *Protoenis*, e.g. in *O. sculda pseudosculda* Korshunov and *Oeneis mongolica* Oberthür (Lukhtanov & Eitschberger, 2000).

Therefore, we suggest to recognize the three genera within this complex: *Oeneis* s.s (excluding *Protoeneis*), *Davidina* s.l. (including *Protoeneis*) and *Neominois.* (1) This system is strongly supported by phylogenetic analysis (Figs 2-4). (2) It is conservative and include only genera that have been always traditionally recognized. (3) The age of the genera corresponds to the criteria described by Talavera *et al*. (2013). (4) In this system, each genus is morphologically homogenous and characterized by at least one clear morphological apomorphy in male genitalia.

### Taxonomic conclusions

As a consequence, we conclude that the name *Protoeneis* Gorbunov, 2001 is congeneric with *Davidina* Oberthür, 1879. The genus *Davidina* consists of the following species: *Davidina diluta* (Lukhtanov, 1994), *D. lederi* (Alpheraky, 1897), *D. mongolica* (Oberthür, 1876), *D. nanna* (Ménétries, 1859), *Davidina sculda* (Eversmann, 1851), *D. tarpeia* (Pallas, 1771), *D. uhleri* (Reakirt, 1866) and *D. urda* (Eversmann, 1847). Thus, *Davidina* is not a local monotypic Chinese endemic genus as it has been previously supposed, but is composed of nine species (Appendix) and has a broad distribution area in the Holarctic region including Europe (where it is represented by *D. tarpeia*) and N. America (where it is represented by *D. uhleri*).

## Acknowledgments

We thank Anatoly Barkalov (Institut of Systematics and Ecology of Animals, Novosibirsk), Blanca Huertas (Natural History Museum, London), Sergei Sinev and Alexander Lvovsky (Zoological Institute of the Russian Academy of Sciences, St. Petersburg), Andrei Sourakov and Andrew Warren (McGuire Center for Lepidoptera and Biodiversity, University of Florida) who provided an opportunity to work with the collections.

The study was supported by the RFBR grant 18-04-00263 to VL and by grant 19-14-00202 from the Russian Science Foundation to the Zoological Institute of the Russian Academy of Sciences. The work of Vladimir Dubatolov was partly supported by the Federal Fundamental Scientific Research Program for 2013-2020, Project No. VI.51.1.5.

## Abbreviations

HT: holotype
LT: lectotype
ST: syntype
TL: type locality
TM: type material

## Museums and institutes

AMNH: American Museum of Natural History, New York, USA.
BMNH: Natural History Museum, London, England [formerly British Museum (Natural History)].
CM: Carnegie Museum of Natural History, Pittsburgh, Pennsylvania, USA.
CNC: Canadian National Collection, Ottawa, Ontario, Canada.
FMNH: Field Museum of Natural History, Chicago, USA.
HUS: Hokkaido University, Sapporo, Japan.
MAKB: Museum Alexander Koenig, Bonn, Germany.
NRS: Naturhistoriska Riksmuseet, Stockholm, Sweden.
SMF: Senckenbergmuseum, Frankfurt am Main, Germany.
SZM: Siberian Zoological Museum, Novosibirsk, Russia.
USNM: National Museum of Natural History, Smithsonian Institution, Washington, D. C., USA.
ZISP: Zoological Institute, St. Petersburg, Russia.
ZMHU: Zoologisches Museum, Humboldt-Universität zu Berlin, Germany.

## Appendix. A catalogue of the genus *Davidina*

### Genus *Davidina* Oberthür, 1879

Et. Entom. **4**: 19, t. 2, fig. 1. Type-species - *Davidina armandi* Oberthür, 1879 (Et. Entom. **4**: 19, t. 2, f. 1).

= *Leechia* Röber, 1907. In Seitz, Grossschmetterlinge der Erde **1**: 43, t. 19b. Type-species - *Leechia alticola* Röber, 1907 (In Seitz, Grossschmetterlinge der Erde 1:43, t. 19b).

= *Sinosatyrus* Lee Chuan-Lung, 1988. Chinese Science Bull. **33** (4): 320. Type-species - *Sinosatyrus beijingensis* Lee, 1988.

= *Protoeneis* P. Gorbunov, 2001. In Gorbunov, The butterflies of Russia: classification, genitalia, keys for identification (Lepidoptera: Hesperioidea and Papilionoidea): 228. Type-species - *Chionobas nanna* Ménétriés, 1859 (Bull. Phys. Acad. St.-Petersb., 3(1): 105).

### 1. *Davidina armandi* Oberthür, 1879

Et. Entom. **4**: 19, t. 2, fig. 1. TL: Pe-hoa-tschan, China. Holotype in BMNH (Fig. 1).

***=*** *alticola* (Röber, 1907) (*Leechia alticola*). In Seitz, Grossschmetterlinge der Erde **1**: 43, t. 19b. TL: “Central-China (Tschang-yang)”. TM in BMNH.

= *beijingensis* (Lee Chuan-Lung, 1988) (*Sinosatyrus beijingensis*). Chinese Science Bull. **33** (4): 320.

**Distribution.** N. E. China.

### 2. *Davidina diluta* (Lukhtanov, 1994)

(*Oeneis diluta*). In Lukhtanov & Lukhtanov, Die Tagfalter Nordwestasiens, Herbipoliana **3**: 138, t. 25, fig. 6. TL: Susch, Ujukski-Gebirge, Tuva, S. Siberia, Russia. HT in ZISP.

**Distribution.** Russia: C. Tuva.

### 3. *Davidina lederi* (Alpheraky, 1897)

(*Oeneis Tarpeia* Pall. var. *Lederi*). Mém. Lép. Rom. **9**: 196. TL: “Urga” [Ulanbator], Mongolia (probably in the Khangai Mts). LT in ZISP, designated by U. Eitschberger & V.A.Lukhtanov (1994), Atalanta **25**:163.

= *sapozhnikovi* Korshunov, 1982 (Oeneis *sapozhnikovi*). Gel’minty, kleshchi i nasekomye: 86, figs 1-2. TL – 10 km NW Mungen-Mort, Central Aimak, Mongolia. HT in SZM.

**Distribution**. N. and C. Mongolia. Russia: Irkut River valley in S. Siberia, S. Tuva.

### 4. *Davidina mongolica* (Oberthür, 1876)

**4a. *Davidina mongolica mongolica* (Oberthür, 1876)** (*Chionobas mongolica*). Et. Entom. **2**: 31, t.4, fig.6. TL: “Mongolie orientale. Elle vole en été dans les montagnes, à une altitude moyenne de 500 mètres” TM in BMNH.

= *tsingtaua* Austaut, 1911 (*Oeneis mongolica tsingtaua*). Ent. Zeitschr. 24: 244. TL: Tsingtau, Shantung, China or. TM - ?.

= *tsingtauana* auct. (misspelling).

= *mandschurica* O. Bang-Haas, 1939 (*Oeneis mongolica mandschurica*). Iris **53**: 49, t.1, fig.6. TL: Kintschou, Prov. Fengtien, Manchuria mer. or., China. TM in ZMHU. f. *extincta* O. Bang-Haas, 1939. Iris **53**:49. TL: Kintschou, Prov. Fengtien, Manchuria mer. or., China. TM in ZMHU.

**Distribution**. N. E. China: Inner Mongolia, Liaoning.

**4b. *Davidina mongolica hoenei* (Groß, 1970**) (*Oeneis mongolica hoenei*). Mitt. Münch. Ent. Ges.

**58**: 22. TL: “Mienshan, Shansi, Nordchina”. HT in MAKB.

**Distribution**. N. China: Shansi [Shanxi].

**4c. *Davidina mongolica coreana* (Matsumura, 1927)** (*Oeneis nanna coreana*). Ins. Mats. **1**: 163, t. 5, fig.9. TL: “Genzan”, Korea. TM in HUS (?).

= *okamotonis* Matsumura, 1927 (“*Oeneis nanna coreana okamotonis* n. ab.” - p. 164; “*Oeneis nanna okamotonis”* - p. 160). Ins. Mats. **1**: 164, t. 5, f.3. TL: “Suigun”, Korea. TM in HUS (?). f. *fumosa* O. Bang-Haas, 1939. Iris **53**:49, t.1, f.6. TL: “Corea int., bei Gensan”. TM in ZMHU.

**Distribution**. N. Korea.

**4d. *Davidina mongolica walkyria* (Fixsen, 1887)** (*Oeneis walkyria*). Mém. Lép. Rom. **3**: 310. t.14, fig.4. TL: “in der Umgebung Pung-Tungs”, Korea. TM in ZISP.

= *shonis* Matsumura, 1927 (“*Oeneis nanna walkyria shonis* n. ab.” - p. 163; “*Oeneis nanna shonis”* - p. 150). Ins. Mats. **1**: 163, t. 5, fig.2. TL: “Mt. Daitoku”, Korea. TM in HUS (?). f. *hakuba* Doi, 1934 (*Oeneis nanna walkyria* f. *hakuba*). In Mori, Doi & Cho, Coloured butterflies from Korea (2): 16, t. 12, fig. 4. TL: Korea. TM - ?. f. *soibona* Doi, 1934 (*Oeneis nanna walkyria* f. *soibona*). In Mori, Doi & Cho, Coloured butterflies from Korea (2): 16, t. 12, fig. 7. TL: Korea. TM - ?.

= ?*masuiana* Matsumura, 1919. Thous. Ins., Add. **3**: 547, t. 38, f. 2. Ins. Mats. **1**:160 (1927). TM-HUS (?).

**Distribution**. C. Korea.

**4e. *Davidina mongolica hallasanensis* (Murayama, 1991)** (*Oeneis urda hallasanensis*). Nature and Insects 26 (3): 20. TL: Mt. Hallasan, Cheju-do Island, S. Korea. TM - ?.

**Distribution**. Korea: Cheju-do Island.

### 5. *Davidina nanna* (Ménétriés, 1859)

**5a. *Davidina nanna dzhulukuli* (Korshunov, 1998)** (*Oeneis anna dzhulukuli*). In Korshunov, Novye opisaniya i dopolneniya dlya knigi “Dnevnye babochki Aziatskoi chasti Rossii”: 27, t. 17(4). TL: Taboshak Mt., Kuraiski Chrebet, Altai, Russia. TM in SZM.

**Distribution**. Russia: S. E. Altai (Kurai, S. Tchuja and Sailjugem Mountains, Ukok-Plateau), W. Tuva.

**5b. *Davidina nanna anna* (Austaut, 1911)** (*Oeneis nanna* var. *anna*). Ent. Zeitschr. **24**: 243. TL: “Arasagun-Gol” (N. Mongolia). TM in ?.

= *brunhilda* A. Bang-Haas, 1912 (*Oeneis brunhilda*). Iris **26**:105, t. 6, f. 1. TL: “aus dem Sajan-Gebiete”. Acording to the syntype labels the type-locality is “Sajan, Arasagun-Gol” (N. Mongolia) and “Sajan, Arsyn” (Russia, S. Tuva). STs in ZMHU.

**Distribution**. Russia: S. Tuva. N. Mongolia.

**5c. *Davidina nanna nanna* (Ménétriés, 1859)** (*Chionobas nanna*). In Schrenk, Reisen und Forschungen im Amur-Lande **2** (1): 38, t. 3, f. 5. TL: “Amour” [Amur], Russia. HT in ZISP.

= *hulda* Staudinger, 1887 (*Oeneis hulda*). Mém. Lép. Rom. **3**: 149, t.16, f.8. TL: Pokroffka, Amur, Russia. LT in ZMHU, designated by F. J. Groß (1970), Mitt. Münch. Ent. Ges. 58: 20, Abb. 29.

= *coriacea* Seitz, 1909 (*Oeneis nanna coriacea*). In Seitz, Grossschmetterlinge der Erde **1**:121, t. 40g. TL: “Apfelgebirge”, S. Siberia, Russia. TM in SMF.

**Distribution**. Russia: Transbaikalia, Amur basin. N E. Mongolia. N.E. China.

**5d. *Davidina nanna jakutski* (Korshunov, 1998)** (*Oeneis jakutski*). Novye opisaniya i dopolneniya dlya knigi “Dnevnye babochki Aziatskoi chasti Rossii”: 27, t. 17(3). TL: Yakutsk, botsad [botanical garden], Yakutia, Russia. TM in SZM.

**Distribution**. Russia: Yakutia.

**5e. *Davidina nanna dzhugdzhuri* (Sheljuzhko, 1929)** (*Oeneis dzhugdzhuri*). Mitt. Münch. Ent. Ges. **19**: 349, t. 28, figs. 3-4. TL: “in der Bergkette Dzgugdzhur (Grenze der Provinzen Amur und Jakutsk), an den Quellen des Flusses Dzhelinda” [Yakutia], Russia. HT in coll. Sheljuzhko (Kiev University, Ukraine).

**Distribution**. Russia: Southern and Eastern Yakutia, northern part of Amurskaya oblast’.

**5f. *Davidina nanna taimyrica* (Lukhtanov & Eitschberger, 2001)**. (*Oeneis nanna taimyrica*). In: Lukhtanov & Eitschberger U. (2001) Catalogue of the genera *Oeneis* and *Davidina* (Bauer E, Frankenbach T, editors) Butterflies of the World, Supplement 4: 33. TL: Russia, Taimyr Peninsula, Putorana plateau, Ayan Lake, near Ayan River. HT in Zoological Museum of Moscow University.

**Distribution**. Russia: Northern Siberia (Taimyr Peninsula).

### 6. *Davidina sculda* (Eversmann, 1851)

**6a. *Davidina sculda sculda* (Eversmann, 1851)** [*Hipparchia* (*Chionobas*) *sculda*]. Bull. Soc. Nat. Mosc. **24**: 612. TL: “environs de Kiachta de la Sibérie orientale” (Buryatia, S. Siberia, Russia). STs in ZISP.

= *velleda* Austaut, 1911 (*Oeneis velleda*). Int. Ent. Zeitschr. **5**: 360. TL: Siberia. TM in ?.

**Distribution**. Russia: Altai, Tuva, Sayan Mts., S. Buryatia. N.E. Mongolia.

**6b. *Davidina sculda pseudosculda* (Korshunov, 1977)** (*Oeneis pseudosculda*). Insects of Mongolia **5**: 662, fig, 1. TL: Sudzykte, Central Aimak, Mongolia. HT in ZISP.

**Distribution**. C. Mongolia.

**6c. *Davidina sculda pumila* (Staudinger, 1892)** (*Oeneis Sculda* Ev. var. *Pumila* Stgr.). Mém. Lép. Rom. **6**: 201. TL: “Pokr.[offka]” and “am oberen Amur” [Transbaikalia region, Russia]. STs in ZMHU.

**Distribution**. Russia: Transbaikal (Tchita region), Amur region. N.E. Mongolia. N.E. China.

**6d. *Davidina sculda vadimi* (Korshunov, 1995)** (*Oeneis sculda vadimi*). In Korshunov & Gorbunov, Dnevnye babochki Asiatskoi chasti Rossii: 138. TL: Severobaikalsk, Verkhneangarski Khrebet, N. Transbaikal, Russia. HT in SZM.

**Distribution**. Russia: N. Transbaikal, Yakutia.

### 7. *Davidina tarpeia* (Pallas, 1771)

**7a. *Davidina tarpeja tarpeia* (Pallas, 1771)** (*Papilio tarpeia*). Reise versch. Prov. russ. Reiches **1**: 171 (*tarpeja*), 480 (*tarpeia*). TL: “In campis aridis ad Volgam copio fus Maio”, Volga region, Russia. TM lost.

*= celimene* (Cramer, 1782) (*Papilio celimene*). Pap. Exot. **4**: 169, t. 375 E, F. TL: “Siberie”.

*= vacuna* Grum-Grshimailo, [1891] (*Oeneis Vacuna*). Horae Soc. Ent. Ross. **25**: 458. TL: “in montibus ad Dongar-tschen”, S. of Xining, Qinghai, China. HT in BMNH.

= *rozhdestvenskyi* Korb et Yakovlev, 1997 (*Oeneis tarpeia rozhdestvenskyi*). Zoosystematica Rossica **5**(2):313. TL: Shtabka, Altai, Russia. HT in ZISP.

= *ukokana* Korb, 1998 (*Oeneis tarpeia ukokana*) Alexanor **20**: 144, figs 1-3. TL: “Südaltai, Ukok, Dschasator”, Russia. HT in ZISP.

= *baueri* Lukhtanov & Eitschberger, 2000 (*Oeneis tarpeia baueri*). Schmetterlinge der Erde **11**: 8. TL: Mezen’ r.[iver], vic. Sush’, Tuva, Siberia, Russia, 1100-1200 m. HT in McGuire Center for Lepidoptera and Biodiversity, University of Florida.

**Distribution**. Russia: E. and S.E. Europe, N. Caucasus (Kislovodsk, Teberda, Elbrus), S. Ural, S.W. Siberia (W. Siberian Lowland), Altai, Tuva. N. and E. Kazakhstan. W. Mongolia. NW. China.

**Note**. There are two different spellings of this name in the original description and in subsequent works. Here we select ***tarpeia*** as the most common spelling in historical and contemporary works.

**7b. *Davidina tarpeia grossi* (Eitschberger & Lukhtanov, 1994)** (*Oeneis tarpeia grossi*). Atalanta **25**: 164, t. Va, figs 1-3. TL: “20 km NW Selenduma”, Buryatia, S. Siberia, Russia. HT in ZISP. **Distribution**. Russia: Buryatia, Chita region. N. Mongolia.

### 8. *Davidina uhleri* (Reakirt, 1866)

**8a. *Davidina uhleri varuna* (W. H. Edwards, 1882)** (*Chionobas varuna*). Canadian Ent. **14**: 205. TL: “plains of Dacotah Terr.”, Canada. LT in CM. ab. *dennisi* Gander, 1927. Canadian Ent. **59**: 285-286 (1927). TL: Beulah, Manitoba. HT in CNC. **Distribution.** Canada: British Columbia, Alberta, Saskatchewan, Manitoba. USA: Montana, Dakotas, Nebraska, W. Minnesota.

**8b. *Davidina uhleri nahanni* (Dyar, 1904)** (*Oeneis nahanni*). Proc. Ent. Soc. Washington **6**:142. TL: Nahanni Mts., Dist. of Mackenzie, Canada. HT in USNM. **Distribution**. Canada: Northwest Territories (Mackenzie Mts).

**8c. *Davidina uhleri cairnesi* (Gibson, 1920)** (*Oeneis cairnesi*). Rept. Canadian Arctic Exped. **3**(1): 15. TL: White River dist., Yukon Territory, Canada. HT in CNC.

=*kluanensis* (Hassler, 2002) (*Oeneis nanna kluanensis*). Nachricht. Entomol. Ver. Apollo **22**: 197-205. TL: SW-Yukon, NW Burwash Landing, Quill Creek Area. HT in SMF. **Distribution**. Canada: Yukon, northwestern corner of the Northwest Territories. USA: N.E. Alaska.

**8d. *Davidina uhleri uhleri* (Reakirt, 1866)** (*Chionobas Uhleri*). Proc. Ent. Soc. Philadelphia **6**:143. TL: “Rocky Mts., Colorado Territory”, USA, restricted to vic. Georgetown, Clear Creek Co., Colorado by F. M. Brown (1953), American Mus. Novitates (1625): 6. TM in FMNH.

= *obscura* (W. H. Edwards, 1892) (*Chionobas Uhleri* var. *Obscura*). Butt. N. America **3**:294. TL: Clear Creek Canyon, Jefferson Co., Colorado. HT in CM.

**Distribution**. USA: Colorado east of the Continental Divide.

**8e. *Davidina uhleri reinthali* (F. M. Brown, 1953)** (*Oeneis uhleri reinthali*). American Mus. Novitates (1625): 10-11. TL: Gothic, Colorado. HT in AMNH.

**Distribution.** USA: western slope of the Continental Divide in Colorado.

### 9. *Davidina urda* (Eversmann, 1847)

**9a. *Davidina urda urda* (Eversmann, 1847)** (*Hipparchia urda*). Bull. Soc. Nat. Mosc. **20**(3): 69, t. 2, f. 1-4.TL: “Dauria”, Russia. TM in ZISP.

= *umbra* Staudinger, 1892 (*Oen. urda* Ev. var. *Umbra*). Iris **5**: 335. TL: “Südamur”, Russia. TM in ZMHU.

*= laeta* Austaut, 1908 (*Oeneis urda* Evers. var. *laeta*). Ent. Zeitschr. **22**: 44. TL: “Sayan”. TM - ?.

= *trybomi* Bryk, 1946 (*Oeneis urda tribomi*). Arkiv Zool. **38A** (3): 25, t. 4, fig. C3. TL: “Krasnojarsk”, [S. Siberia, Russia]. HT in NRS. ab. *albidior* Austaut, 1908. Ent. Zeitschr. **22**: 44. TL: “Sayan”. TM - ? ab. *banghaasi* Austaut, 1908. Ent. Zeitschr. **22**: 44. TL: “Sayan”. TM -?

**Distribution**. Russia: N. Altai (Tongosh Mountains), S. Siberia, Transbaikal, Amur and Primorye regions, S. and Central Yakutia. N. Mongolia. N.E. China.

**9b. *Davidina urda tschiliensis* (O. Bang-Haas, 1933)** (*Oeneis urda tschiliensis*). Ent. Zeitschr. **47**: 98. TL: Hsingan mont., Tunkia-jingze, Prov. Tschili, China sept. HT is probably in ZMHU. **Distribution**. N.E. China: Hebei.

**9c. *Davidina urda monteviri* Bryk, 1946** (*Oeneis urda monteviri*). Arkiv Zool. **38A** (3): 24, t. 4, fig. A3. TL: “Shinten”, Korea. HT in NRS(?).

**Distribution**. N. Korea.

## Notes

### Competing Interest Statement

The authors have declared no competing interest.

